# A niche-derived non-ribosomal peptide triggers planarian sexual development

**DOI:** 10.1101/2023.12.06.570471

**Authors:** Melanie Issigonis, Katherine L. Browder, Rui Chen, James J. Collins, Phillip A. Newmark

## Abstract

Germ cells are regulated by local microenvironments (niches), which secrete instructive cues. Conserved developmental signaling molecules act as niche-derived regulatory factors, yet other types of niche signals remain to be identified. Single-cell RNA-sequencing of sexual planarians revealed niche cells expressing a non-ribosomal peptide synthetase (*nrps*). Inhibiting *nrps* led to loss of female reproductive organs and testis hyperplasia. Mass spectrometry detected the dipeptide β-alanyl-tryptamine (BATT), which is associated with reproductive system development and requires *nrps* and a monoamine-transmitter-synthetic enzyme (AADC) for its production. Exogenous BATT rescued the reproductive defects after *nrps* or *aadc* inhibition, restoring fertility. Thus, a non-ribosomal, monoamine-derived peptide provided by niche cells acts as a critical signal to trigger planarian reproductive development. These findings reveal an unexpected function for monoamines in niche-germ cell signaling. Furthermore, given the recently reported role for BATT as a male-derived factor required for reproductive maturation of female schistosomes, these results have important implications for the evolution of parasitic flatworms and suggest a potential role for non-ribosomal peptides as signaling molecules in other organisms.

## Introduction

Biogenic monoamines (including serotonin, dopamine, histamine, and tryptamine) act as important neurotransmitters and neuromodulators across the animal kingdom. However, many of these “neurotransmitters” are not exclusive to animals: they are also found in plants, unicellular organisms, and even bacteria. Therefore, monoamines predate the existence of nervous systems and have non-neuronal functions, which are not well characterized (1, 2).

Monoamine synthesis is catalyzed by a widely conserved enzyme, Aromatic L-amino acid decarboxylase (AADC). Recent work uncovered an unanticipated role for AADC in germ cell development and regeneration in the sexual strain of the planarian *Schmidtea mediterranea* (3). In addition to the previously reported expression of *aadc* in planarian serotonergic and dopaminergic neurons as well as photoreceptor pigment cells (4–6), *aadc* expression was enriched in the somatic gonadal “niche” cells of ovaries and testes, and in vitellaria, the yolk cell-producing reproductive organs. RNA interference (RNAi)-mediated inhibition of *aadc* in sexual planarians resulted in ovary ablation and loss of accessory reproductive organs, including vitellaria, oviducts, sperm ducts, and the gonopore. Knockdown of *aadc* also resulted in hyperplastic testes, with a dramatic expansion of germline stem cells and lack of differentiated male germ cells (3). The striking phenotypes observed in ovaries, testes, and accessory reproductive organs, combined with the enriched expression of *aadc* in niche cells of the reproductive organs, suggested that these tissues may be responding to a localized monoamine niche signal from within the gonads themselves. However, the systemic nature of RNAi in planarians (7–9) and the importance of the nervous system for their reproductive development (10–13), meant that roles for neurally derived monoamines could not be ruled out.

## Results

### *nrps* expression in somatic gonadal cells

To investigate the source of the monoamine signal and characterize the transcriptomes of somatic gonadal niche cells, we performed single-cell RNA sequencing (scRNA-seq) on sexual planarians, focusing on regions of the animal containing reproductive organs (Fig. 1A). Recent scRNA-seq analyses of *S. mediterranea*, although highly informative, focused on the asexual strain, which reproduces by transverse fission and lacks mature gonads and accessory reproductive organs; thus, these studies did not define cell types comprising the reproductive organs (14, 15). Examination of the sexual planarian single-cell data revealed that *aadc* was highly expressed in a cluster of 73 cells that co-expressed three known somatic gonadal markers: the regulator of male identity *dmd1* (16), the orphan GPCR *ophis* (11), and the extracellular matrix component *LamA* (17) (Fig. 1B and Fig. S1, A to C). Among the top “cluster-defining” genes expressed exclusively in this group of cells is SMESG000023215.1 (Fold change=9.8, p-value=6.3^−21^) (Fig. 1B), a gene encoding a predicted non-ribosomal peptide synthetase (NRPS) similar to *Drosophila ebony* (18) and *Schistosoma mansoni nrps* (*Sm-nrps*)(19) (Fig. S2).

**Fig 1.**
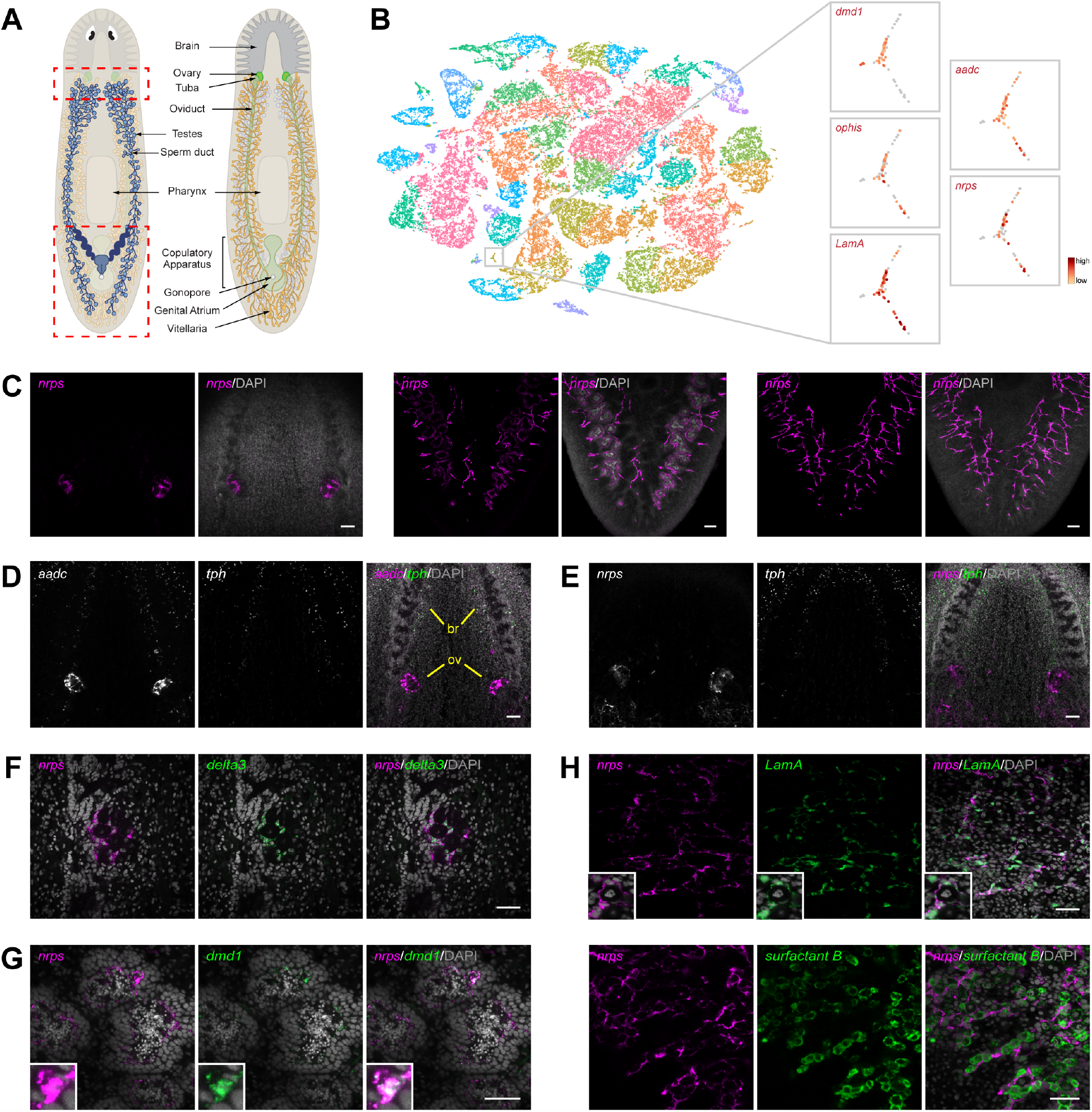
*nrps* is expressed in somatic niche cells in gonads and vitellaria. (**A**) Schematic of dorsal (left) and ventral (right) views of various reproductive organs and landmark structures in adult sexual *S. mediterranea*. (**B**) tSNE plot of 48,216 cells from sexual *S. mediterranea* clustered by shared gene expression into 39 major clusters. t-SNE plots of 73 cells that are highly enriched (red) for somatic gonadal gene markers *dmd1, ophis, LamA*, and *aadc*. Cells in this cluster also express *nrps*. (**C**) Maximum-intensity projections of confocal sections of *nrps* FISH (magenta) showing ovarian expression in the ventral head region (left), testes in the dorsal tail region (middle), and vitellaria in the ventral tail region (right). Note the presence of branches from the vitellaria surrounding the testes in the middle panels. (**D** and **E**) Projection showing dFISH of *aadc* or *nrps* (white/magenta) and serotonergic marker *tph* (white/green) in ventral head region. *aadc*, but not *nrps*, is expressed in *tph*^*+*^ serotonergic neurons. *aadc* and *nrps* are expressed in the ovary (ov) at the base of the brain (br). (**F**) Single confocal section of *nrps* (magenta) and *delta3* (green) co-expression in ovary. (**G**) dFISH of *nrps* (magenta) and somatic niche marker *dmd1* (green) in testes. Insets are high-magnification views of *nrps*^*+*^ *dmd1*^*+*^ cell body. (**H**) dFISH of *nrps* (magenta) with *LamA* (green), or yolk cell marker *surfactant b* (green) in vitellaria of sexually mature planarians. Insets are high-magnification views of *nrps*^*+*^ *LamA*^*+*^ somatic support cells surrounding a yolk cell. Nuclei are counterstained with DAPI (gray; C-H). Scale bars, 100 μm (C-E), 50 μm (F-H).

To validate the single-cell sequencing results, we used fluorescent in situ hybridization (FISH) to examine *Smed-nrps* (for brevity, referred to hereafter as *nrps*) expression in mature sexual planarians. We detected *nrps* expression exclusively in ovaries, testes, and vitellaria (Fig. 1C). Unlike *aadc*, which is also expressed neuronally with the serotonergic marker *tryptophan hydroxylase* (*tph*; Fig. 1D), *nrps* expression is restricted to the gonads and vitellaria (Fig. 1, C and E). To define the gonadal cell type(s) in which *nrps* is expressed, we performed double FISH to detect *nrps* and known somatic gonadal markers. In the ovaries, *nrps* expression is restricted to the *delta3*^*+*^ and *aadc*^*+*^ somatic gonadal cells that envelop the oogonia and oocytes (Fig. 1F and Fig. S3A). Similarly, in the testes, *nrps* is expressed in *dmd1*^*+*^ and *ophis*^*+*^ somatic gonadal cells, which form long cytoplasmic projections that encyst the developing male germline (Fig. 1G and Fig. S3B). In vitellaria, *nrps* is expressed in the *LamA*^*+*^ *aadc*^*+*^ somatic support cells that envelop *surfactant b*^*+*^ yolk cells (12) (Fig. 1H and Fig. S3C). Thus, *nrps* is expressed in niche cells in female and male reproductive organs.

### *nrps* RNAi phenocopies *aadc* knockdown

Having demonstrated that *nrps* is a niche cell marker in gonads and vitellaria, we asked whether it is required for development of these reproductive organs. We performed *nrps* RNAi by feeding hatchlings double-stranded RNA (dsRNA) twice a week for 4 weeks (the amount of time it takes for control planarians to reach sexual maturity). We tested for *nrps* RNAi specificity by feeding animals dsRNA targeting three distinct regions of *nrps* (Fig. S4A) and confirmed by quantitative PCR that *nrps* was significantly knocked down (Fig. S4B). We examined the effects of *nrps* knockdown on female reproductive development by double FISH to detect the germline stem cell marker *klf4l* (17) with early germ cell marker *nanos* (17, 20–23) or somatic niche marker *ophis. nrps* RNAi resulted in a significant reduction or complete loss of *klf4l*^*+*^ *nanos*^*+*^ germ cells in the anterior ovarian fields located mediolaterally along the brain and within the ovaries themselves. All ovaries were devoid of mature germ cells (Fig. 2A). Similarly, *nrps* RNAi animals did not have any vitellaria (Fig. 2B). The ablation of female germ cell and yolk cell lineages in *nrps* RNAi animals likely results from loss of *ophis*^*+*^ ovarian somatic gonadal cells and vitelline support cells (Fig. 2, B and C); in the absence of their respective niches, female germ and yolk cells are not maintained.

**Fig. 2.**
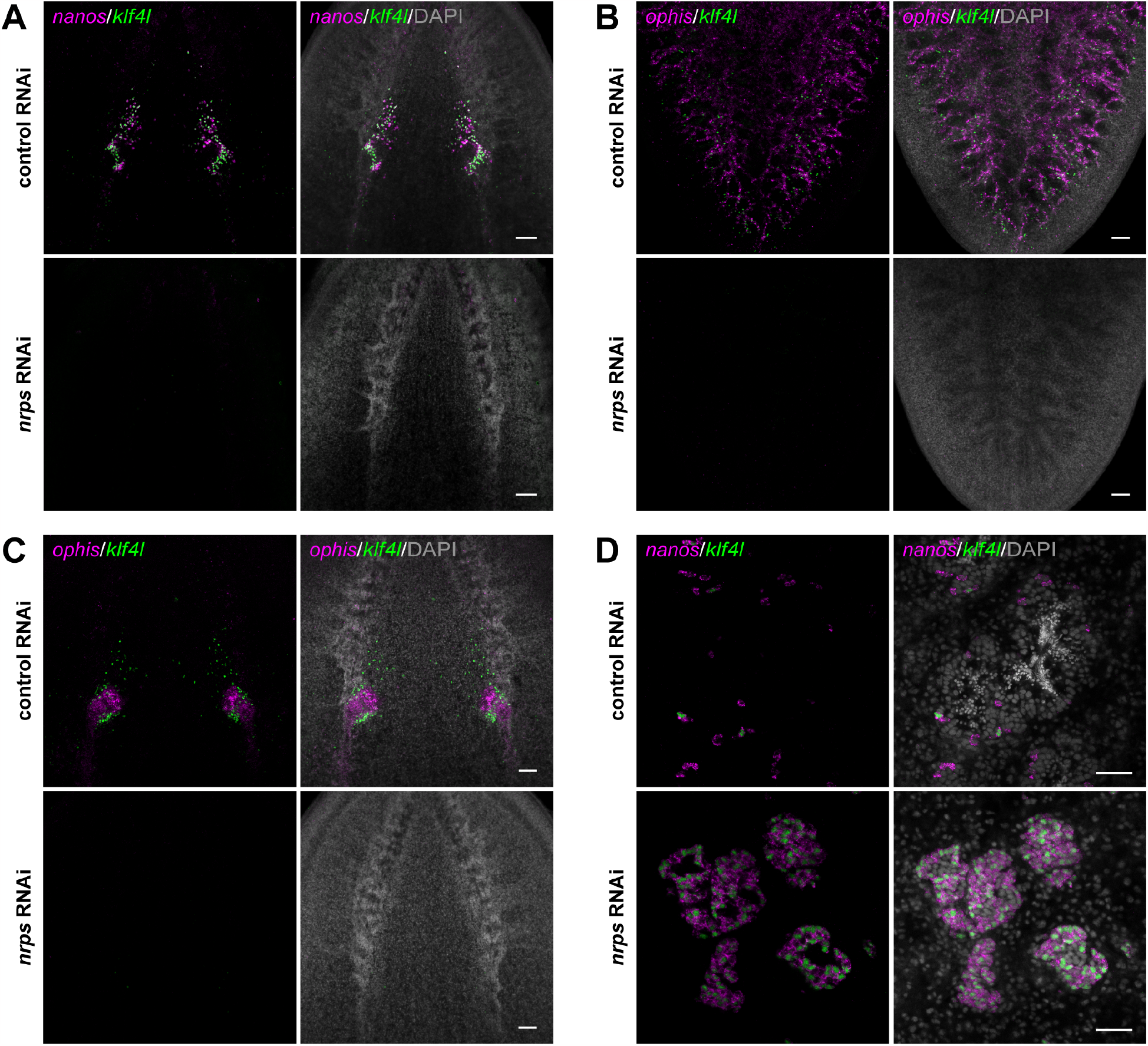
*nrps* is required for gametogenesis, vitellogenesis, and maintenance of somatic niche cells in adult ovaries and vitellaria. (**A**) Projections of confocal sections showing dFISH of *klf4l* (green) with *nanos* (magenta) in the ovarian fields and ovaries located posterior to the brain of sexually mature planarians. (**B**) Projections of confocal sections showing dFISH of *klf4l* (green) with *ophis* (magenta) in the vitellaria in the ventral posterior region. (**C**) Projections of confocal sections showing dFISH of *klf4l* (green) with *ophis* (magenta) in the ovaries. (**D**) Confocal sections showing dFISH of *klf4l* (green) and *nanos* (magenta) in testes of sexually mature control vs *nrps* RNAi planarians. *nrps* RNAi results in hyperplastic testes consisting of excess *klf4l*^*+*^ *nanos*^*+*^ early germ cells and lacking sperm. Nuclei are counterstained with DAPI (gray; A-D). n=12-15 animals per RNAi condition. Scale bars, 100 μm (A-C), 50 μm (D).

In contrast to the loss of ovaries, hyperplastic testes develop in *nrps* RNAi animals. In control testes, early *klf4l*^*+*^ *nanos*^*+*^ germ cells are found around the periphery, with spermatogenesis progressing towards the lumen where mature sperm are released. By contrast, *nrps* RNAi testes displayed a dramatic expansion of early *klf4l*^*+*^ and *nanos*^*+*^ germ cells and completely lacked differentiated male germ cells (Fig. 2D). The phenotypes observed in *nrps* RNAi animals – loss of female reproductive tissues and testis hyperplasia –are similar to those found in *aadc* knockdowns.

### *nrps* and *aadc* are required for the synthesis of the dipeptide β-alanyl-tryptamine

The similarities in their expression patterns and the phenotypes observed in *aadc* and *nrps* knockdown animals suggested that these enzymes may operate in the same pathway. Support for this idea comes from the reported functions of NRPS in other animals. *ebony* is a well-known *Drosophila* gene: fly mutants displaying a darkly pigmented, or “ebony,” cuticle color were first described over a century ago (24). Molecular cloning of *ebony* revealed that it encodes a non-ribosomal peptide synthetase that conjugates β-alanine to monoamine transmitters, like dopamine and histamine, to generate β-alanyl-monoamine conjugates (18, 25). More recently, the product of a related *nrps* from the parasitic flatworm *Schistosoma mansoni* was shown to conjugate β-alanine to tryptamine, generating the dipeptide β-alanyl-tryptamine (BATT)(19). In contrast to the modification catalyzed by *Drosophila* Ebony, which inactivates monoamine signals, the production of BATT by the male schistosome triggers reproductive maturation of his female partner (19).

To identify the product(s) of NRPS activity in *S. mediterranea*, we performed liquid chromatography-mass spectrometry (LC-MS) analysis on whole planarian extracts (Fig. 3A). Of the possible β-alanyl-monoamine conjugates, we only detected BATT in planarians and found that mature sexual planarians synthesize significantly more BATT than either sexual hatchlings or asexual planarians, both of which lack mature reproductive organs (Fig. 3A). BATT was undetectable following knockdown of either *nrps* or *aadc*, whereas BATT abundance was unaffected by dsRNAs targeting control bacterial sequences or a paralogous *nrps* gene expressed in the gut (Fig. 3B). Furthermore, BATT abundance was significantly decreased after *ophis* RNAi, which causes defects in the somatic niches of ovaries, testes, and vitellaria, but not in *klf4l* knockdowns, which affect the germ cell and yolk cell lineages. Thus, production of BATT requires the functions of both *nrps* and *aadc*. These data are consistent with a pathway in which AADC produces the monoamine tryptamine from tryptophan and then NRPS catalyzes the conjugation of β-alanine to tryptamine, to generate BATT in somatic gonadal niche cells (Fig. 3C).

**Fig. 3.**
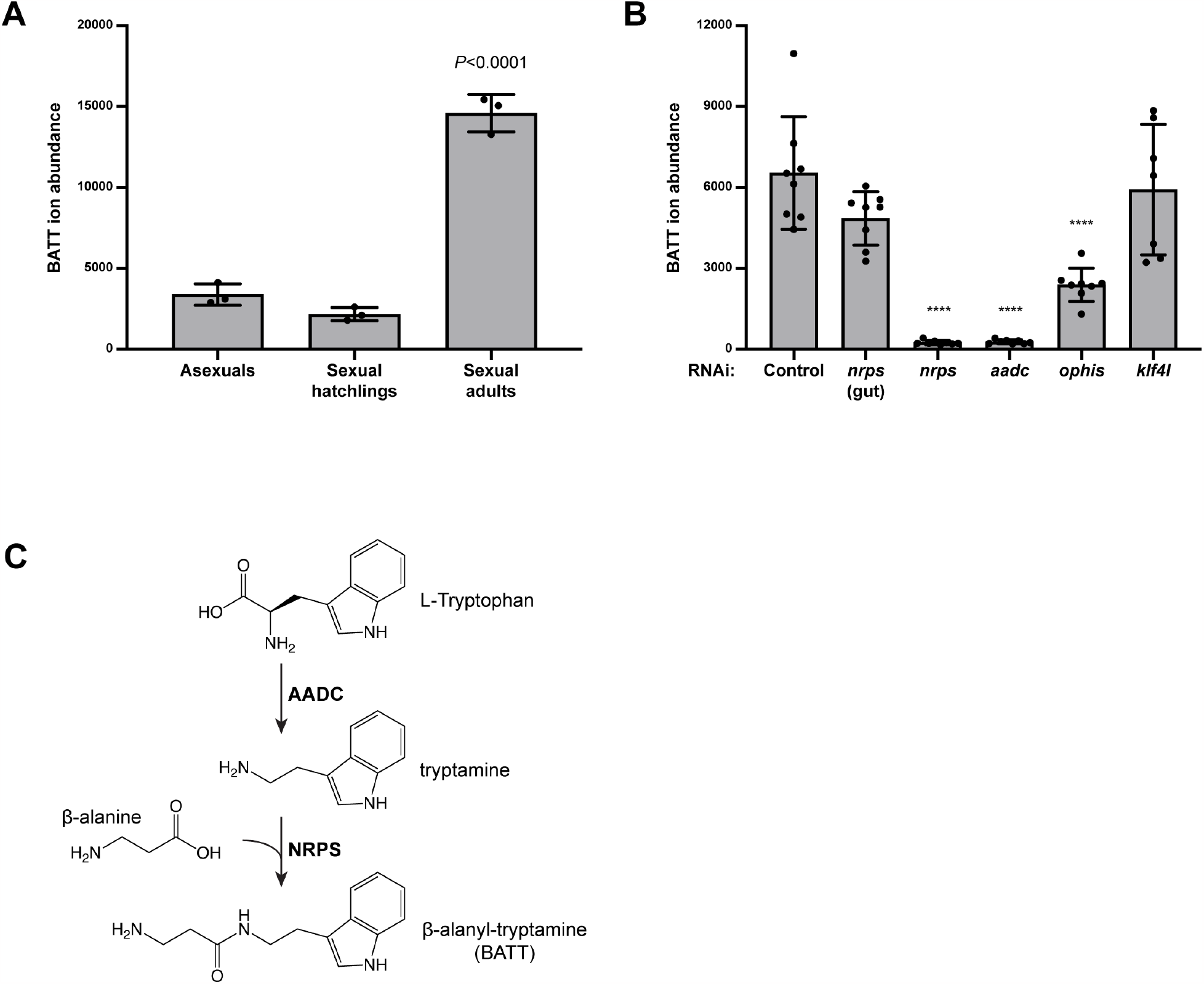
BATT synthesis in sexual planarians requires *nrps* and *aadc* in somatic gonadal niche cells. (**A**) Relative abundance of BATT ions detected by LC-MS in asexual planarians, sexually immature hatchlings, and sexually mature adults. Mature sexual planarians have significantly more BATT compared to either asexual animals or sexual hatchlings both of which lack reproductive structures. n=3 biological replicates, *p* < 0.0001, one-way ANOVA. Data are presented as mean ± SD. (**B**), Relative abundance of BATT ions in sexually mature planarians after the following RNAi treatments: control, *nrps* (gut; a paralogue of *nrps* expressed in the gut but not in reproductive tissues acts as a specificity control), *nrps, aadc, ophis* (somatic niche cells in the ovaries, testes, and vitellaria), *klf4l* (female/male germline stem cells and yolk cell progenitors). n=8 biological replicates per RNAi condition. *p* < 0.0001, one-way ANOVA. Data are presented as mean ± SD. (**C**) Proposed pathway for BATT synthesis: AADC catalyzes decarboxylation of L-tryptophan to the monoamine tryptamine, and subsequently NRPS conjugates β-alanine to tryptamine to produce the dipeptide BATT.

### BATT rescues *nrps* and *aadc* knockdowns

If the dramatic phenotypes observed following RNAi-mediated knockdown of *nrps* or *aadc* result from an inability to synthesize BATT, then supplying exogenous BATT to knockdown animals should restore development of their reproductive system. Increasing concentrations of chemically synthesized BATT were delivered to planarians via feeding. Control RNAi animals exposed to synthetic BATT (0.1-100 μM) all displayed wild-type ovaries. In contrast, the ovary-loss phenotype observed in *nrps* knockdown planarians was rescued in a dose-dependent manner when they were exposed to increasing concentrations of synthetic BATT (Fig. 4A). With 100 μM BATT supplementation, ovaries were indistinguishable from controls. Intriguingly, wild-type planarians receiving the highest dose of BATT displayed precocious sexual maturation, as indicated by the appearance of a gonopore (Fig. S5).

**Fig. 4.**
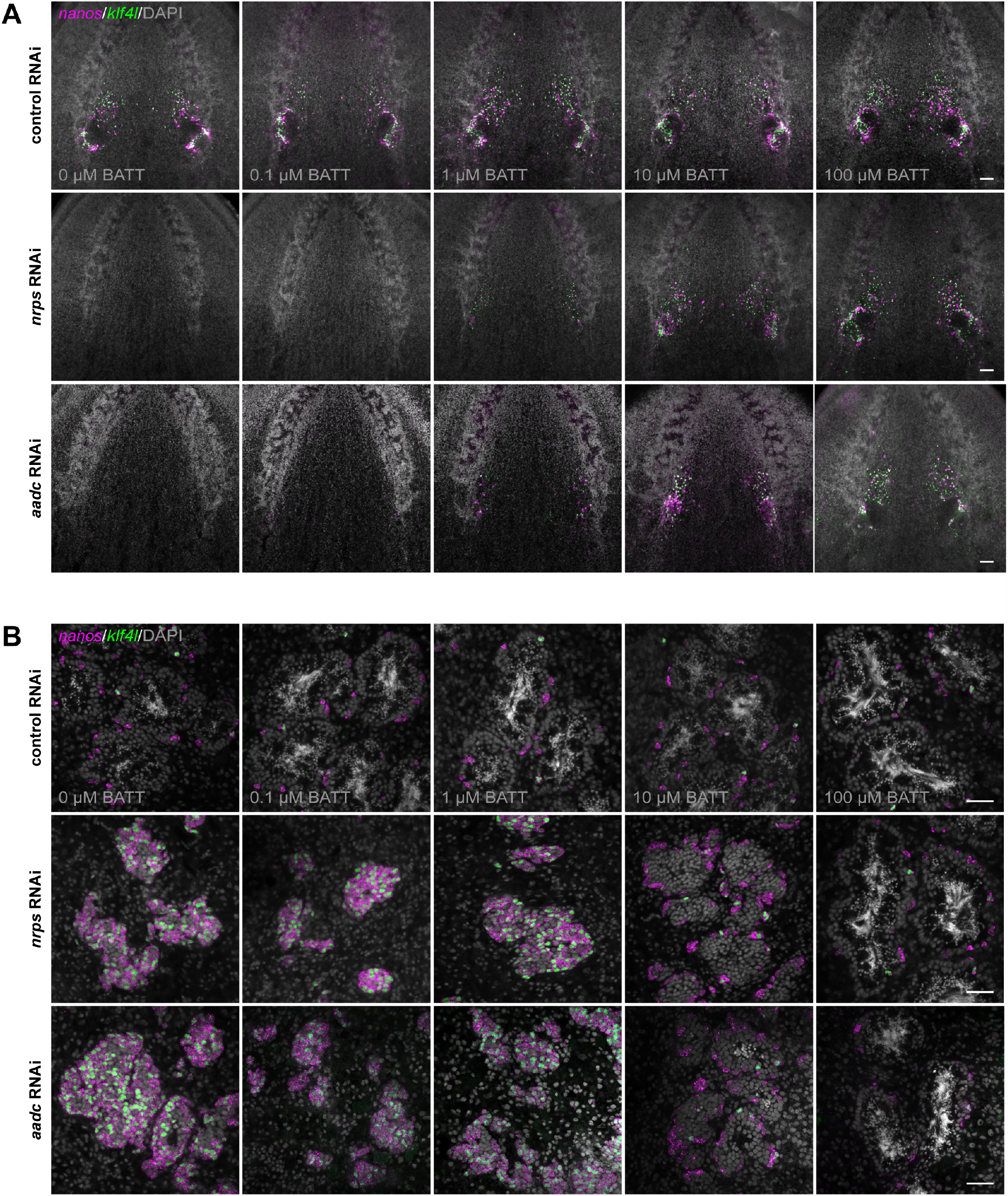
BATT rescues *nrps* and *aadc* knockdowns. (**A**) Projections of confocal sections showing dFISH of *klf4l* (germline stem cells; green) and *nanos* (early germ cells; magenta) in the ventral head region of adult sexual RNAi animals. RNAi animals were supplemented with indicated concentrations of BATT by feeding. Ovaries from control RNAi animals fed increasing concentrations of BATT appear wild type; *klf4l*- and *nanos*-expressing cells are found along the periphery of the ovaries and in anterior ovarian fields. Ovary ablation in *nrps* and *aadc* knockdowns is rescued in a dose-dependent manner by BATT feeding; ovaries are indistinguishable from controls at the highest BATT concentration (100 μM). (**B**) Confocal sections of *klf4l* (green) and *nanos* (magenta) dFISH showing few *klf4l*/*nanos* double-positive germline stem cells and *nanos* single-positive early germ cells along each testis periphery. All stages of spermatogenesis are present in control animals +/–BATT. The testis hyperplasia phenotype observed in *nrps* and *aadc* knockdowns is rescued by BATT supplementation. Nuclei are counterstained with DAPI (gray; A and B). N=3 experiments, n=7-21 planarians per RNAi condition. Scale bars, 100 μm (A), 50 μm (B).

Unlike control testes, which contain a handful of *klf4l*^*+*^ *nanos*^*+*^ germline stem cells around the periphery, all stages of spermatogenesis, and mature sperm in the lumen, testes of *nrps* RNAi animals that were not exposed to BATT were hyperplastic and filled primarily with *klf4l*^*+*^and *nanos*^*+*^ undifferentiated germ cells. However, when exposed to increasing concentrations of BATT, testes from *nrps* knockdown animals were rescued back to wildtype in a dose-dependent manner (Fig. 4B). Importantly, beyond morphological rescue after BATT exposure, reproductive function was fully restored: these animals laid eggs that produced viable embryos. Exposure of *aadc* RNAi planarians to exogenous BATT resulted in similar restoration to wild-type ovary and testis function (Fig. 4, A and B). Thus, the reproductive system defects of *aadc* RNAi or *nrps* RNAi planarians are due to loss of the BATT signal.

## Discussion

Here we showed that the non-ribosomal peptide BATT acts as a critical signal to activate reproductive system development in the planarian (Fig. 5). Because the expression of *nrps*, which is required for BATT synthesis, is restricted to somatic niche cells, these cells appear to be the source of this signaling molecule. The expression of the monoamine transmitter biosynthetic enzyme AADC in *nrps*^*+*^ somatic niche cells, the similarity between *aadc* and *nrps* knockdown phenotypes, as well as the rescue of these knockdowns by exogenous BATT reflect a non-neuronal role for AADC: production of tryptamine as a substrate for β-alanylation by NRPS, ultimately yielding BATT.

**Fig. 5.**
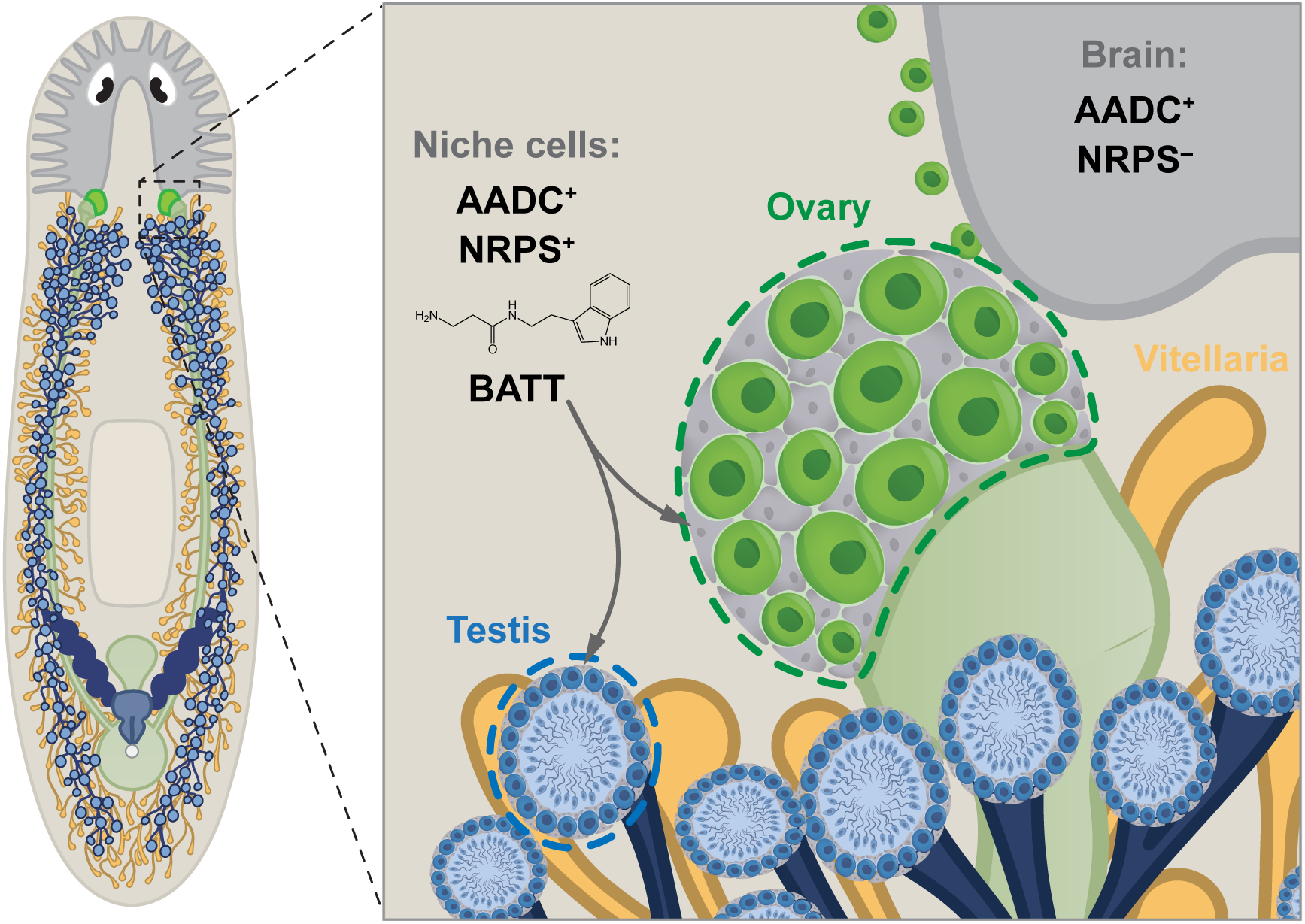
Niche cells produce the monoamine-derived dipeptide β-alanyl-tryptamine to regulate germ cell development. Model showing that in addition to the conserved role of AADC in serotonergic and dopaminergic neurons in the nervous system, it produces the monoamine tryptamine in the somatic niche cells of various reproductive organs that co-express NRPS. Niche cells that co-express AADC and NRPS produce β-alanyl-tryptamine/BATT, which acts as a localized niche signal regulating germ cell development.

Based on analysis of *Drosophila* Ebony, β-alanylation of monoamines was thought simply to inactivate these important signaling molecules. For example, in *Drosophila* epidermal cells, Ebony converts the melanin precursor dopamine to β-alanyl-dopamine (26–28). In *ebony* mutants, excess dopamine accumulates and is converted into melanin, resulting in hyperpigmented flies (29). In glial cells surrounding the photoreceptors, Ebony converts the photoreceptor neurotransmitter histamine to β-alanyl-histamine (carcinine), inactivating it and facilitating its recycling back to photoreceptors (18, 25, 30–32). *ebony* mutants display optomotor and visual system defects(33–35), revealing the importance of this mode of regulating dopamine and histamine metabolism.

The recent identification of BATT as a male-derived trigger of schistosome female reproductive development (19) revealed that such β-alanyl-monoamine conjugates can also act as signals in their own rights. Finding that planarians also deploy BATT as a critical reproductive signal demonstrates that the use of such β-alanyl-monoamine conjugates as signaling molecules is not a parasite-specific innovation. How are these novel signals transduced? Do they signal through G-protein coupled receptors (GPCRs) like related monoamines, or do other players mediate their signaling activities? The opposite effects of *nrps* knockdown on female and male germ cell development (loss of ovaries, development of hyperplastic testes) provide a useful system for dissecting the mechanisms downstream of the BATT signal.

In schistosome males, expression of *nrps* is triggered in a ventral population of neurons upon pairing with the female partner; release of BATT from the male then induces female sexual development and egg laying (19). The expression of BATT synthetic machinery in somatic niche cells of planarians provides a striking contrast with the male-specific, neuronal expression observed in schistosomes. Is this alteration in the source of BATT in schistosomes relative to the free-living planarian related to their parasitism or the evolution of separate sexes (which is unique among flatworms)? Have male schistosomes been converted into the functional equivalent of a somatic niche for their partners? Wider sampling of *nrps* expression and BATT function throughout the flatworms (both free-living and parasitic) should help resolve these questions.

NRPS enzymes are typically found (and best characterized) in bacteria and fungi, in which they generate diverse metabolites ranging from virulence factors and toxins to antibiotics (36). Aside from a handful of examples (37–39), NRPS function in metazoans represents largely uncharted territory. The observation that both planarians and schistosomes use BATT as a reproductive signal suggests an ancient and conserved role for *nrps* genes, at least among flatworms. Because *nrps* related sequences have been identified in many other animals (39), future work may reveal novel monoamine conjugates acting as signaling molecules more broadly throughout the animal kingdom.

## Materials and Methods

### Planarian culture

Sexual *S. mediterranea* (40) were maintained in 0.75X Montjuïc salts (41) at 18°C. Asexual *S. mediterranea* (clonal strain CIW4) (42) were maintained in 1X Montjuïc salts at 21°C. Planarians were reared on a calf liver puree diet and starved for 1 week before experimentation.

### Single-cell RNA sequencing

Single-cell suspensions were generated as previously described (14) from tissues enriched for reproductive organs (outlined in Fig. 1a) from sexually mature *S. mediterranea*. Briefly, tissue sections from 15-20 planarians were finely diced into small fragments (<1 mm) and transferred to a 15 ml conical with CMFB (Calcium-Magnesium Free solution; 400 mg/L NaH_2_PO_4_, 800 mg/L NaCl, 1200 mg/L KCl, 800 mg/L NaHCO_3_, 1.33 mM glucose, 15 mM HEPES pH 7.3, 1% BSA) supplemented with 10 μg/ml collagenase. Cell dissociations were assisted by mechanical agitation on a nutator with intermittent pipetting. Cells were dissociated for ∼10 minutes, passed through a 40-μm filter, and pelleted by centrifugation for 5 minutes at 310 x *g*. After supernatant removal, the cell pellet was resuspended in 4 ml of CMFB. Cells were incubated in Hoechst 33342 (10 mg/mL) for 45 minutes and then stained with propidium iodide (PI; 10 μg/μL) immediately before sorting on a FACSAria (BD Biosciences). Single, live cells were collected and processed for 10X Genomics sequencing as described (43). Sequencing reads were aligned to the sexual *S. mediterranea* S2F2 genome (version SMESG.1) (44, 45) using Cell Ranger 3.0 (10X Genomics). Dimensionality was reduced in Loupe Browser (10X Genomics) and visualized with graph-based clustering using t-distributed stochastic neighbor embedding (t-SNE).

### Cloning genes for riboprobe or dsRNA synthesis

Cloning was performed as previously described (10). Briefly, genes (Table S1) were cloned by PCR amplification of cDNA synthesized from RNA isolated from sexual *S. mediterranea*. Gene-specific primers (Table S2) were used and amplified products were ligated into pJC53.2 plasmid via TA cloning. Anti-sense riboprobes for in situ hybridization experiments were generated by in vitro transcription reactions with T3 RNA polymerase (46). dsRNA for RNAi experiments was generated by in vitro transcription reactions with T7 RNA polymerase (9).

### In situ hybridization

FISH protocols were performed as previously described (46, 47) with the following modifications for sexual planarians (17): Adult sexual planarians were killed in 7.5% N-acetyl-L-cysteine in 1X PBS for 10 minutes; fixed in 4% formaldehyde in PBSTx (1X PBS + 0.1% Triton X-100) for 25-30 minutes; bleached in Bleaching Solution (1X SSC solution containing 5% deionized formamide and 1.2% hydrogen peroxide) for 4 hours; incubated in PBSTx containing 10 μg/ml Proteinase K and 0.1% SDS for 20 minutes; and refixed in 4% formaldehyde in PBSTx for 15 minutes. Planarians were blocked in Blocking Solution (5% heat inactivated horse serum, 5% Roche Western Blocking Buffer in TNTx [0.1 M Tris pH 7.5, 0.15 M NaCl, 0.3% Triton X-100]) for 2 hours at room temperature, and incubated in Blocking Solution containing anti-Digoxigenin-POD (1:2,000) or anti-Dinitrophenyl-HRP (1:2,000) for 8 hours at 12°C. For fluorescent development of riboprobes, TSA reactions were performed for 30 minutes.

### RNA interference

dsRNA was generated as previously described (9), diluted in a final volume of 80-100 μl of water, and annealed. dsRNA was mixed with homogenized calf liver (1:3) and 1-2 μl of food coloring and fed to animals two times per week for 4 weeks. dsRNA generated from the bacterial *CamR* and *ccdB*-containing insert of pJC53.2 was used for control RNAi feedings (10).

### BATT supplementation

BATT (MuseChem) (19), β-alanine (Sigma-Aldrich, 146064), and tryptamine (Sigma-Aldrich, 193747) were added to planarian food (liver, or liver and dsRNA) at final concentrations of 100, 10, 1, or 0.1 μM.

### Imaging

Confocal imaging was performed with a ZEISS LSM 880 or 980 with a C-Apochromat 10x/0.45 W M27 or Plan-Apochromat 40x/1.3 Oil DIC M27 objective. Linear adjustments and maximum intensity projections were made with ZEISS ZEN 3.1 (Blue) software.

### Liquid chromatography-mass spectrometry (LC-MS)

To detect endogenous levels of BATT, planarians were flash-frozen in liquid Nitrogen and homogenized in 1 mL ice-cold 80% methanol in water (vol/vol) using an ultrasonicator (30 seconds × 3 times at the highest setting). Homogenates were vortexed and 200 μL was diluted in 800 μL ice-cold 80% methanol. This mixture was vortexed again for 1 minute, centrifuged at maximum speed for 15 minutes at 4°C, and this supernatant (1 mL) containing the metabolome extract was transferred to new tube and stored at -80°C. To normalize between samples, protein concentrations in sample pellets were quantified using a BCA assay (Thermo Scientific, 23225). LC/MS studies using metabolome extracts (corresponding to 70 or 100 μg of protein, depending on the replicate) were performed as previously described (19).

### Quantitative PCR

Quantitative PCR (qPCR) was performed as previously described (17). Briefly, total RNA was extracted with TRIzol reagent from RNAi planarians that had been starved 1 week after the final dsRNA feeding. Total RNA was DNase-treated, purified (RNA Clean & Concentrator Kit, Zymo Research Corporation), and reverse transcribed into cDNA (iScript cDNA Synthesis Kit, Bio-Rad Laboratories, Inc.). *nrps* primers (Table S2) were designed to sequences outside of the dsRNA target sequence (i.e., the sequence cloned into pJC53.2). qPCR reactions were set up with GoTaq qPCR Master Mix reagent system (Promega) and run on a StepOnePlus Real-Time PCR System (Applied Biosystems, Thermo Fisher Scientific). Gene expression levels were normalized to the endogenous control gene *β-tubulin*. The ΔΔCt method was used to analyze relative changes in *nrps* expression in RNAi knockdowns (4 biological replicates, 3 technical replicates each). 2^−ΔΔCt^ values (normalized to control RNAi) with 95% confidence intervals were calculated in Microsoft Excel and plotted using GraphPad Prism software.

### Protein Alignment

MUSCLE alignment of the following protein sequences was performed using Jalview software (using default settings): *Drosophila melanogaster* Ebony (NP_524431.2), *S. mediterrranea* NRPS, *Schistosoma mansoni* NRPS (A0A3Q0KR05.1).

## Acknowledgments

We thank Newmark/Issigonis lab members, past and present, for discussion and feedback. We especially thank Akshada Redkar for excellent technical support and help with planarian cell dissociations, and John Brubacher and Tania Rozario for invaluable comments on the manuscript. The authors used the University of Wisconsin – Madison Biotechnology Center Gene Expression Center (Research Resource Identifier – RRID:SCR_017757) for single-cell RNA sequencing. This work was supported by NIH R01 HD043403 (PAN) and R01 AI150776 (JJC), and Welch Foundation award I-1948-20180324 (JJC). PAN is an investigator of the Howard Hughes Medical Institute.

