## Supplementary material for "A niche-derived non-ribosomal peptide triggers planarian sexual development": Fig. S

**Supplemental Figures and Tables**

#### Fig. S1

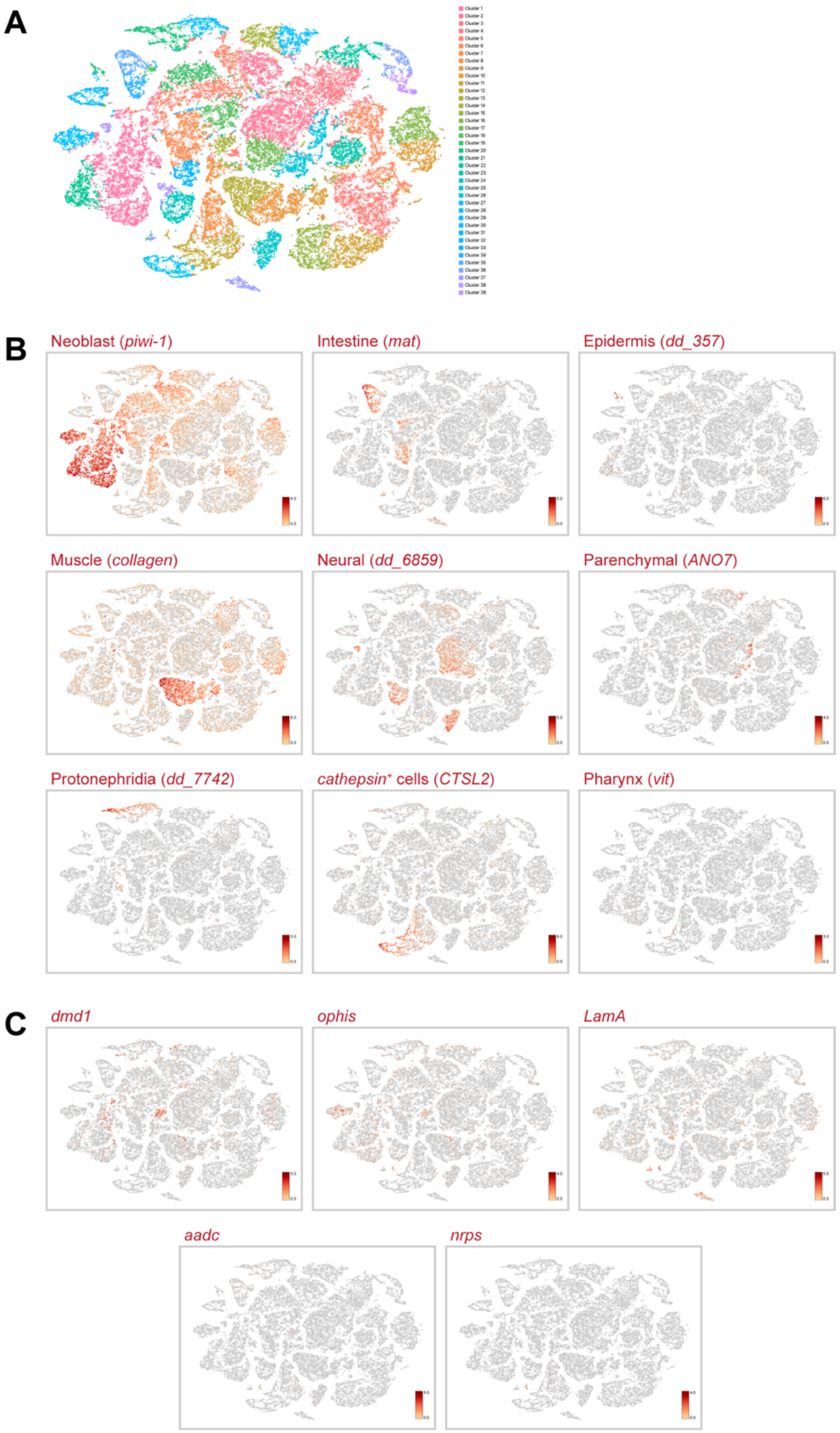

**Fig. S1. Single-cell RNA sequencing of cells from sexual *S. mediterranea*.** (**A**) tSNE plot of 39 clusters. (**B**) t-SNE plots of representative genes for nine major planarian tissue classes previously characterized in asexual *S. mediterranea* (14). All somatic tissue classes are present except for pharyngeal cells, which are not detected in this sexual sc-seq dataset since we enriched for reproductive tissues lacking this organ. (**C**) t-SNE plots showing expression of somatic gonadal gene markers *dmd1*, *ophis*, *LamA*, *aadc*, and *nrps*.

### Fig. S2

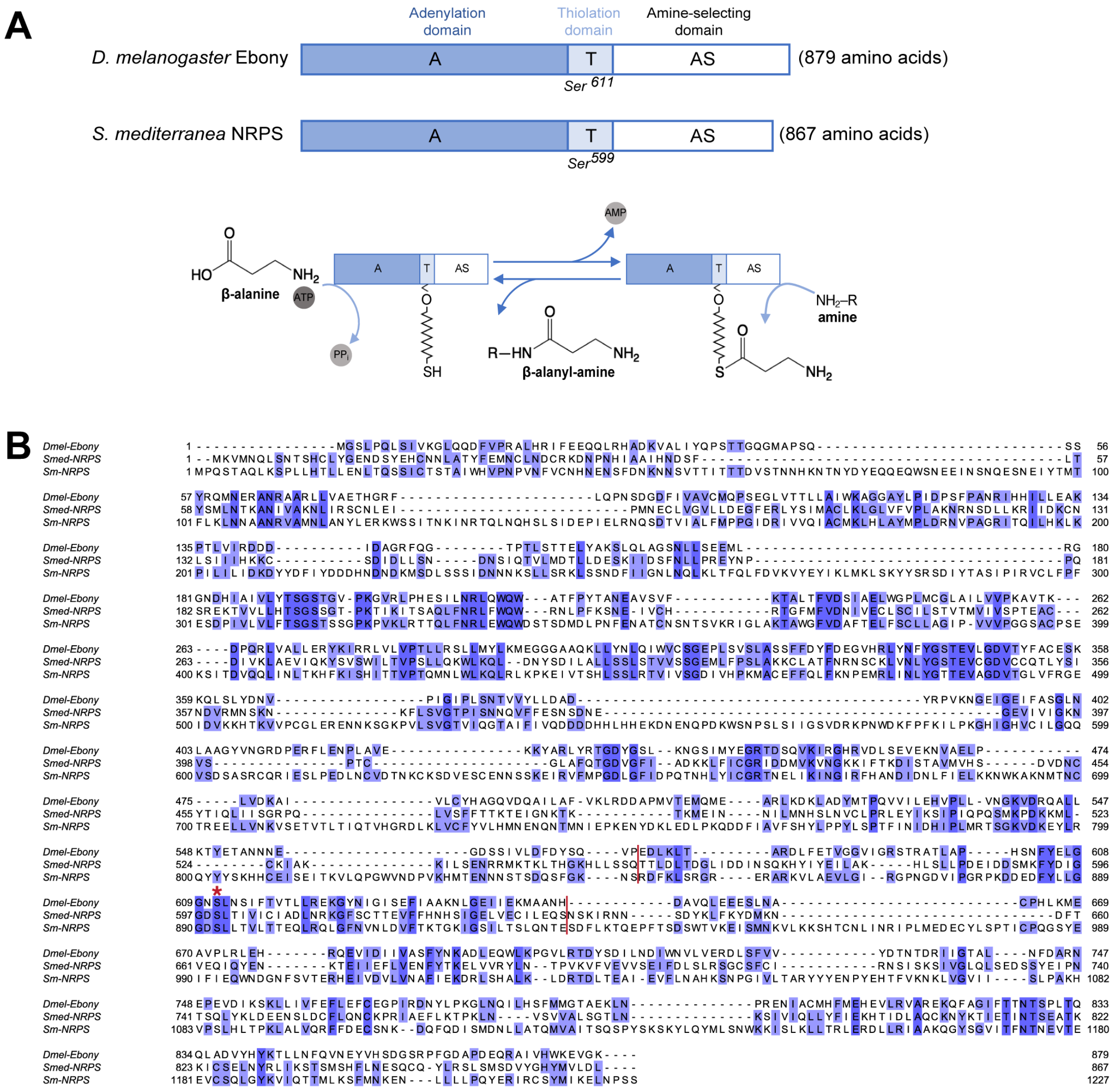

**Fig. S2. *Smed-nrps* encodes a non-ribosomal peptide synthetase.** (**A**) Schematic showing adenylation (A), thiolation (T), and amine-selecting (AS) domains in *D. melanogaster* Ebony and the *Schmidtea mediterranea* homolog NRPS. *Drosophila* and *S. mediterranea* share a conserved serine residue in their thiolation domains. NRPS proteins like Ebony can conjugate β-alanine to various biogenic amines (e.g., dopamine, histamine, etc.). This enzymatic process involves three steps: adenylation of β-alanine catalyzed by the adenylation (A) domain; covalent attachment of β-alanine to a phosphopantetheinyl group on a conserved serine within the thiolation (T) domain; and binding of an amine in the amine-selecting (AS) domain, which facilitates nucleophilic attack of the NRPS-bound β-alanine resulting in a β-alanyl-amine dipeptide product. (**B**) Protein alignment of *Drosophila* Ebony, *S. mediterranea* NRPS, and *S. mansoni* NRPS. The serine thiolation site (marked by an asterisk) in the T domain (delineated by red lines) is conserved.

### Fig. S3

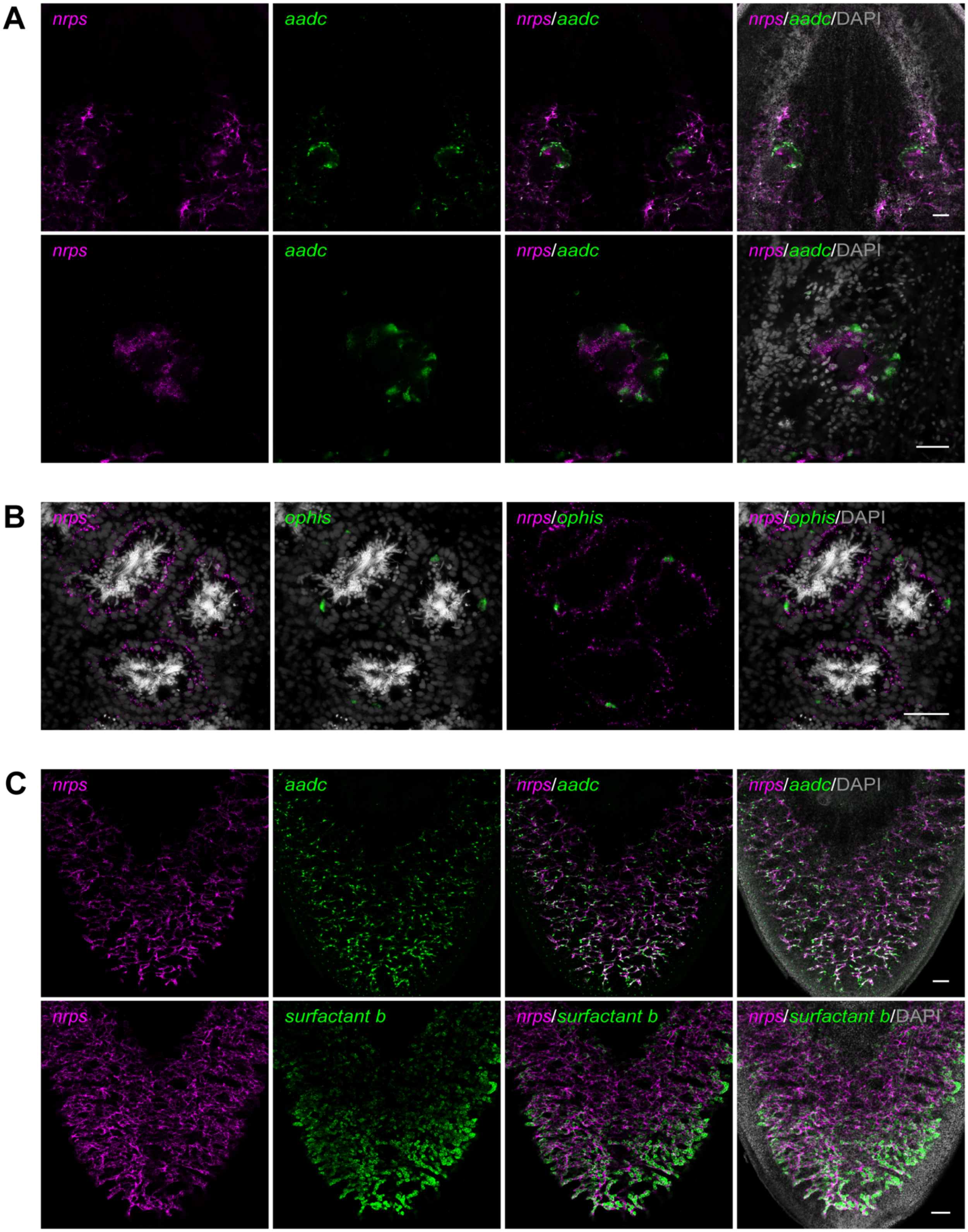
A
nrps
aadc
nrps/aadc
nrps/aadc/DAPI
B
nrps
ophis
nrps/ophis
nrps/ophis/DAPI
C
nrps
aadc
nrps/aadc
nrps/aadc/DAPI
surfactant b
nrps/surfactant b
nrps/surfactant b/DAPI

**Fig. S3. *nrps* is expressed in somatic gonadal niche cells.** (**A**) Projection of ventral head region (top) and confocal section of ovary (bottom) showing dFISH of *nrps* (magenta) and *aadc* (green). (**B**) Confocal section of testes with *nrps* (magenta; cytoplasmic localization) and *ophis* (green; nuclear) co-expressing cells. *ophis* RNA localizes mainly to the nucleus of somatic gonadal cells, which extend long *nrps*$^{+}$ cytoplasmic projections that encyst developing germ cells. (**C**) Projections of confocal sections showing dFISH of *nrps* (magenta) with *aadc* (green; top), or yolk cell marker *surfactant b* (green; bottom) in the ventral posterior region of sexually mature planarians. Nuclei are counterstained with DAPI (gray; A-C). Scale bars, 100 µm (A, top; C), 50 µm (A, bottom; B).

### Fig. S4

#### A *nrps* (SMEST023215002.1):

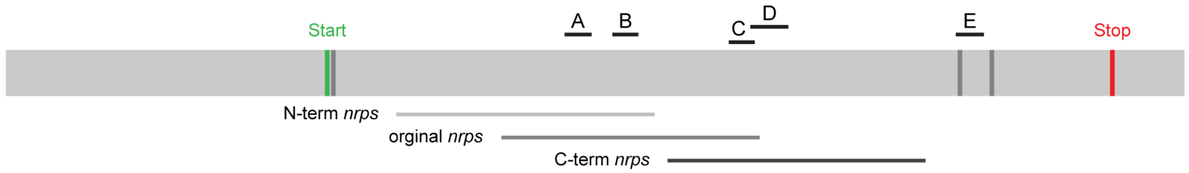

## B

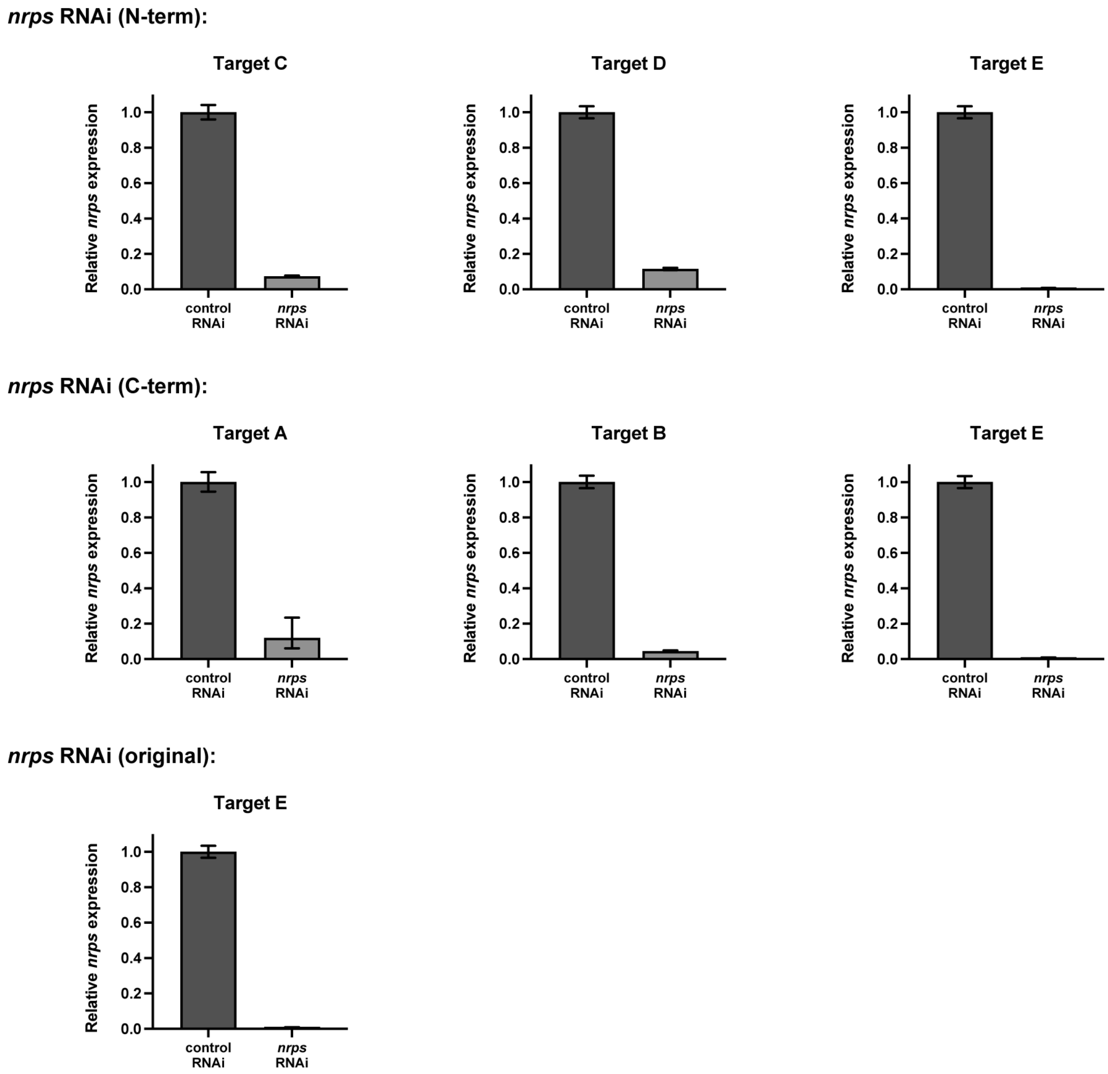

**Fig. S4. Testing *nrps* RNAi specificity and quantifying *nrps* expression in knockdown animals.** (**A**) To test for *nrps* RNAi specificity and exclude the possibility of off-target effects, RNAi was performed with dsRNA targeting three ~1 kb regions of *nrps*: the original region used throughout this study, and 2 non-overlapping regions (N-terminus vs C-terminus). *nrps* gene (gray bar) is shown with positions for start (green) and stop (red) codons, exon-exon boundaries (dark gray), cloned regions (bottom), and qPCR amplicons (top: A-E). (**B**) qPCR analysis of *nrps* mRNA expression normalized to *β-tubulin* in control and *nrps* RNAi animals depicting efficient knockdown of *nrps* after RNAi. Top: dsRNA targeting the N-terminus of *nrps* was used for RNAi-mediated knockdown of *nrps*, and qPCR primers targeting regions C, D, and E were used to quantify *nrps* expression levels. Middle: dsRNA targeting the C-terminus of *nrps* was used for RNAi and qPCR primers targeting regions A, B, and E were used to quantify *nrps* expression levels. Bottom: dsRNA targeting the original cloned amplicon of *nrps* and qPCR primers targeting region E were used to quantify *nrps* expression levels. N = 4 biological replicates (3 technical replicates each). Bar graphs depict relative quantification ($2^{-\Delta\Delta Ct}$) values normalized to control RNAi with 95% confidence intervals.

#### Fig. S5

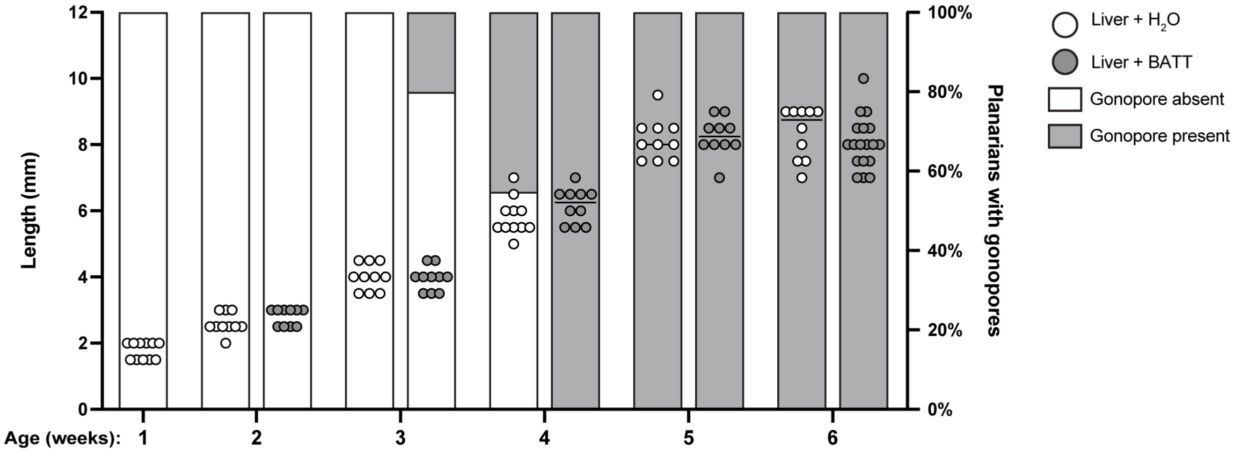

**Fig. S5. BATT triggers sexual maturation.** Quantification of sexual planarian length (mm; left Y axis; horizontal line represents median) and gonopore presence (right Y axis) during development. One-week old hatchlings were fed liver +/–BATT for 6 weeks. Supplementation with BATT did not affect growth but triggered precocious sexual maturation (evidenced by the presence of a gonopore) in +BATT individuals. n=10-18 planarians per time point.

**Table S1. Information for transcripts mentioned in this paper.**

| Gene | Reference | Sequence information |
|---|---|---|
| ***nrps*** | This paper | Planmine SMEST.1: SMEST023215002.1 |
| ***nrps*** **(gut)** | This paper | Planmine SMEST.1: SMESG000017098.1 |
| ***klf4l*** | Issigonis et al., 2022 | Planmine SMEST.1: SMEST031008001.1 |
| ***Laminin A*** | Issigonis et al., 2022 | Planmine SMEST.1: SMEST056013009.1 |
| ***delta3*** | Khan et al., 2022 | Genbank accession: OL957299 |
| ***nanos*** | Wang et al., 2007 | Genbank accession: EF035555.1 |
| ***dmd1*** | Chong et al., 2013 | Genbank accession: KC736555.1 |
| ***ophis*** | Saberi et al., 2016 | Genbank accession: KX018822.1 |
| ***surfactant B*** | Rouhana et al., 2017 | Genbank accession: KY847536.1 |
| ***tph*** | Curie & Pearson, 2013 | Genbank accession: KF134114.1 |
| ***aadc*** | Curie & Pearson, 2013 | Planmine SMEST.1: SMEST022173002.1 |

**Table S2. Primer sequences.**

| Cloning into pJC53.2 | Forward | Reverse |
|---|---|---|
| ***nrps*** | TCGTTTGCCACAGAACAGGA | CGATTAGGCCGTCGGTTAGG |
| ***nrps* (gut)** | CCGATTTTGGCAGCTTCTGG | CTCAGCCACCCATTTCGTCT |
| ***nrps* (N-term)** | TGGATGAAGGCTTTGAGAGG | GATTCGGCCGCAAATAAATA |
| ***nrps* (C-term)** | ACAGCCGTTATGGTCCACTC | ACCGACACGTTTAGCTTTGG |
| **qPCR** | **Forward** | **Reverse** |
| ***β-tubulin*** | TGGCTGCTTGTGATCCAAGA | AAATTGCCGCAACAGTCAAATA |
| **Target A** | TCGTAAGCAGCGGTGAAAT | CTTCCGTTGAGCCGTAAAGA |
| **Target B** | GCACACCCATTTCGAACAATC | CAGTCCACAAGTCGGTGATAC |
| **Target C** | TGCTCTGTAAGATTGCGAAGA | GTCTAGAGTGGTTTGGGATGAG |
| **Target D** | CCCAAACCACTCTAGACCTAAC | GAGTCACCACCAATATCGTAGAA |
| **Target E** | ACGTCTGAAGCAACGAAGAA | GTGAGCGCAAATATTGACATTGA |
